# PLAID: ultrafast single-sample gene set enrichment scoring

**DOI:** 10.1101/2025.06.14.659661

**Authors:** Antonino Zito, Xavier Escribà Montagut, Gabriela Scorici, Axel Martinelli, Murodzhon Akhmedov, Ivo Kwee

## Abstract

**Motivation:** In recent years, computational methods have emerged that calculate enrichment of gene signatures within individual samples, rather than across pooled samples. These signatures offer critical insights into the coordinated activity of functionally related genes, proteins or metabolites, enabling identification of unique molecular profiles based on gene set and pathway activity in individual cells and patients. This strategy is pivotal for patient stratification and advancement of personalized medicine. However, the rise of large-scale datasets, including single-cell profiles and population biobanks, has exposed significant computational inefficiencies in existing methods. Current methods often demand excessive runtime and memory resources, becoming impractical for large datasets. Overcoming these limitations is a focus of current efforts by bioinformatics teams in academia and the pharmaceutical industry, as essential to support basic and clinical biomedical research.

**Results:** To address this critical need, we developed PLAID (<u>P</u>athway <u>L</u>evel <u>A</u>verage <u>I</u>ntensity <u>D</u>etection), an ultrafast and memory optimized single sample gene set enrichment algorithm that utilizes sparse matrix computation. PLAID delivers highly accurate gene set scoring and surpasses the performance of current methods in single-cell and bulk transcriptomics, and proteomics data. A distinctive feature of PLAID is its integration of the most widely used gene set scoring algorithms, enabling researchers to apply multiple methods for cross-validation with outstanding runtime efficiency and minimal memory requirement.

**Availability and Implementation:** PLAID is implemented in the R language for statistical computing. PLAID source code and installation instructions are available with no restrictions at https://github.com/bigomics/plaid

## 1 Introduction

Single-sample -including single cell-resolution analyses are rapidly expanding our understanding of biology [Haque et al., 2017; Van de Sande et al., 2023; Zheng et al., 2025]. Single-cell transcriptomics and epigenomics profiles enable identification of disease-related molecular signatures and inter-individual differences, supporting development of personalized medicine [Sheridan, 2024; Pekayvaz et al., 2025]. While single-gene studies have uncovered individual dysregulated factors, complex diseases typically involve heterogenous pathophysiology, driven by multi-factorial perturbations in cellular networks. As these networks mediate homeostatic regulations, minor dysregulation within a gene set might impact critical cellular functions [Galindez et al., 2013; Yang 2020]. Hence, targeted therapeutics -including gene restoration-are most effective when they restore the function of the entire gene set or pathway, not just single genes. Characterizing these pathways enable patient stratification and personalized treatments [Fröhlich et al., 2018].

Curated collections such as MSigDB [Liberzon et al., 2015] and Reactome [Milacic et al., 2024], serve as reference for gene set scoring. Gene set enrichment methods map genes to these sets and compute ‘enrichment scores (ESs)’, whose statistical significance can be assessed using competitive or self-contained null hypothesis tests [Goeman et al., 2007]. While earlier methods, such as GSEA [Subramanian et al., 2005], scored the gene sets at population level, newer methods score gene sets within each sample or cell (Barbie et al., 2009; Hänzelmann et al., 2013; Aibar et al., 2017; Foroutan et al., 2018; Pont et al., 2019; Andreatta et al., 2021]. Using individual sample ranks as basis for ESs bypasses the need to recalculate ESs when integrating new samples. Furthermore, earlier approaches were mostly supervised, modeling ESs on discrete or continuous traits. Differently, unsupervised strategies, such as ssGSEA [Barbie et al., 2009], assess coordinately down-or up-regulated feature signatures by relying on relative feature ranking rather than absolute expression. Another method, Gene Set Variation Analysis (GSVA) [Hänzelman et al., 2013], leverages the global samples’ distribution to assess low or high gene expression in each sample, normalized to a Gaussian or discrete Poisson distribution.

A common drawback among most current single-sample gene set ES methods is the computational inefficiency as sample size increases. This is especially true with the rise of large single-cell data. Both ssGSEA and GSVA are impractical in terms of runtime and memory usage. For instance, ssGSEA requires over 600 seconds to process 2,864 gene sets across 1K single cells on a modern multi-core, 48GB RAM laptop. GSVA, though sometimes faster, is often even more demanding in memory (Fig.1). To address these limitations, more efficient approaches have been developed. Single-cell Signature Explorer (scSE) computes a score for each single cell transcriptome and a given gene set as the sum of all UMI for the genes in the gene set, divided by the total UMI in the cell. Relying on sum -which is computationally efficient- and developed in Go language, scSE is capable to compute 13 millions scores (1K gene sets; 13K cells) in under 5 minutes, outperforming several approaches [Pont et al., 2019]. Ucell takes a different approach, scoring gene signatures within a sample using the Matt-Whitney U statistics [Andreatta et al., 2021; Mann et al., 1947]. Specifically, genes in each cell are ranked and a pseudo-rank of 1 added to mitigate zero-inflated counts. Despite these advances, the rapid growth of population-scale and single-cell data—sometimes with hundreds of thousands of cells—means that computational inefficiency remains a major barrier. Development of scalable, memory-efficient algorithms is essential to prevent bioinformatics bottlenecks and advance precision medicine and systems biology.

**Figure 1.**
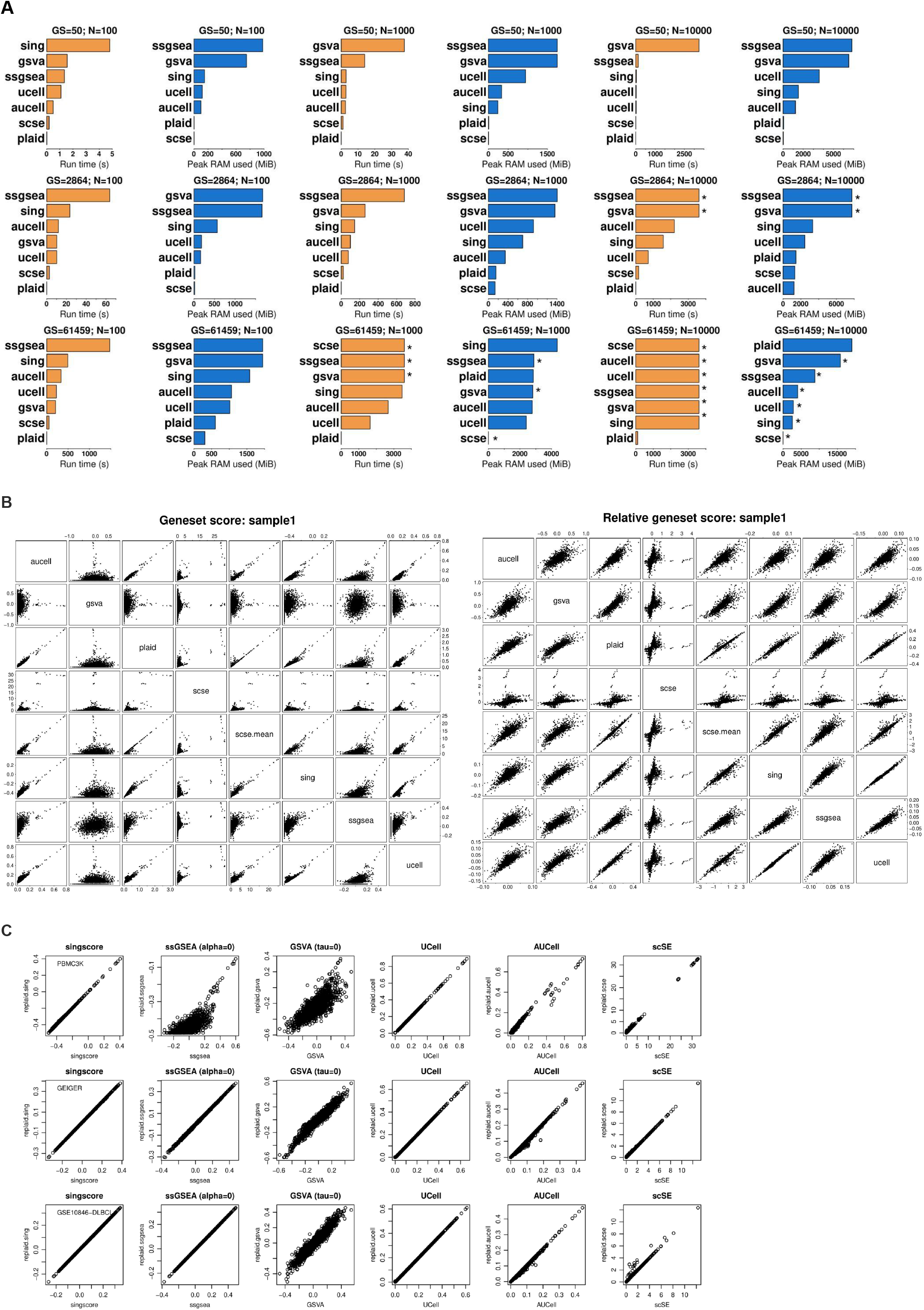
Benchmarking PLAID runtime, memory, and enrichment scores against singscore, ssGSEA, GSVA, Ucell, AUCell, and scSE. The PBMC3K single-cell RNA-seq (scRNAseq) 10X Genomics dataset, including ∼2700 single cells and available from the Seurat R package [Hao et al., 2023]. **(A)** Run time (seconds) and peak RAM memory (MiB) usage. Memory usage was calculated using the PeakRAM R package [Quinn, 2017]. Benchmarking was conducted on data and gene sets of increasing sizes: 100 cells, 1K cells, 10K cells (this latter generated by replication of cells up to 10K cells); 50 gene sets, corresponding to the OmicsPlayground collection of ‘HALLMARK’ gene sets [Akhmedov et al., 2020]; 2864 gene sets, corresponding to the OmicsPlayground collection of ‘GO_BP’ gene sets; 61459 gene sets, corresponding to the whole OmicsPlayground collection of gene sets. Runs were timed out at 1h (3600s). For timed out runs, the real, effective peak memory usage could not be correctly estimated. Timed out runs are indicated by an asterisk. **(B)** Pairwise scatter plots between PLAID vs. singscore, ssGSEA, GSVA, UCell, AUCell and scSE and scSE.mean ESs (Methods). On the left, original ESs. On the right, row-centered gene set ESs. 2864 GO_BP gene sets were used for analysis. **(C)** Pairwise scatter plots between original and PLAID-replicated ESs for different methods (left to right): singscore, ssGSEA, GSVA, UCell, AUCell and scSE. Three testing datasets were used: (top to bottom): PBMC3K scRNAseq dataset; LC/MS proteomics dataset (Wolf et al., 2020); mRNA microarray (GSE10846; Lenz et al., 2008). In most cases, the PLAID (re)-implementation of the distinct enrichment methods provides accurate replication, with the key advantage of being significantly faster and memory efficient.

To address the pressing need for scalable and efficient gene set ES methods, we developed PLAID (<u>P</u>athway <u>L</u>evel <u>A</u>verage Intensity <u>D</u>etection). PLAID scores each gene set based on the average intensity of its constituent genes within each sample. PLAID stands out as a highly-performing solution for single-sample gene set ES. It is ultrafast and memory efficient due to optimized use of sparse matrices. PLAID delivers ESs that are highly concordant with existing methods, while achieving up to 100-fold gain in computational efficiency. We conducted extensive testing of PLAID using single-cell and bulk transcriptomics, and proteomics data. PLAID reliably identifies enriched gene sets, with ESs concordant with other leading methods. Notably, PLAID maintains its speed and memory efficiency in datasets with large sample sizes, outperforming the widely used singscore [Foroutan et al., 2018], ssGSEA [Barbie et al., 2009], GSVA [Hänzelmann et al., 2013], UCell [Andreatta et al., 2021] and AUCell [Aibar et al., 2017]. These methods become impractical due to excessive runtimes and memory requirements. Validating significantly enriched gene sets through independent methods remains critical for robust analysis, however, current methods may hinder this best practice for large data. To provide a solution to this computational barrier, we’ve implemented a unified framework in which singscore, ssGSEA, GSVA, scSE, UCell and AUCell are available within PLAID and use PLAID as ‘back-end’. This architecture enables researchers to perform cross-method validation workflows while maintaining PLAID’s signature speed and memory efficiency. Ultimately, this is particularly useful for gene set scoring in large single-cell and bulk datasets.

Altogether, we show that PLAID not only matches established methods, but provides unmatched computational efficiency, making it uniquely suited for large-scale applications. PLAID represents a significant advancement in single-sample gene set scoring, combining accuracy, scalability, and reproducibility with remarkable speed and resource efficiency.

## 2 Methods

### PLAID algorithm

PLAID requires: (i) log-transformed data (gene, protein, metabolite expression), with features on rows and samples on columns, herein referred to as *X*. Ideally, *X* is a sparse matrix; (ii) a sparse gene set matrix with features on rows and gene sets on columns, herein referred to as *G*. Often, gene sets are available as gmt (Gene Matrix Transposed), which can be converted into a sparse matrix using the provided *gmt2mat* function. Both *X* and *G* are subsetted to include shared features. The sum of each column in *G* is computed, corresponding to the number of genes present in each gene set, and added with a 1e-8 offset to prevent full zeros. Each column is scaled by its sum using the Matrix::colScale R function to handle sparse matrices. The normalization of *G* ensures that columns sum to 1. PLAID single-sample ES (PLAID ‘*ssES*’) is calculated using the matrix cross-product between *G* and *X*, as follows:

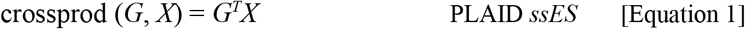

equivalent to the R call t(*G*) %*% *X*. Mathematically, the formula above is the core computation of PLAID, corresponding to the average log-intensity of all features within a gene set in a sample. For large number of gene sets (e.g., >10K) or cells (e.g., >100K), PLAID calculates the cross-product in chunks to avoid memory burden. PLAID does not center nor ranks the input matrix. It outputs the results as a score matrix, with gene sets on rows and samples on columns.

### Normalization of PLAID gene set enrichment scores

The output of PLAID is a matrix with gene sets on rows and samples on columns. By default, PLAID applies median normalization of the results. Specifically, each column is median-centered (so that median is 0). The overall mean of the column medians is then added. Because gene set enrichment scores are normalized, it is not strictly necessary to normalize the input matrix *X*. The median-normalized gene set matrix can be used for differential gene set expression testing between groups.

### Sparse matrix computations

PLAID significantly relies on sparse matrix computation. This is possible with the Matrix R package [Maechler et al., 2000], which provides functions for operating on sparse matrices, wrapping the ‘SuiteSparse’ libraries [Davis et al]. In fact, single cell data are preferably represented as sparse matrices. It is highly recommended to use sparse matrices whenever possible as this will take advantage of the superior computational efficiency for gene set scoring by PLAID. We’ve developed the helper functions *gmt2mat* and *mat2gmt* to convert a gmt gene set list to a sparse matrix and vice versa. Creating the sparse matrix from a gmt of 61,459 gene sets took about 47.5 seconds (Linux AMD Ryzen 4750U, 8-core, 48GB. Ubuntu 24.04.2 LTS), resulting in a matrix with >99% sparsity. Once prepared, the gene set matrix was reused throughout analyses.

### PLAID as a suite of other single-sample gene set enrichment scoring methods

PLAID efficiency is largely attributed to use of sparse matrices. While conducting our benchmarking analyses against existing methods, we also aimed at improving existing methods’ efficiency. We’ve implemented functionalities to use singscore [Foroutan et al., 2018], GSVA [Hänzelmann et al., 2013], UCell [Andreatta et al., 2021], AUCell [Aibar et al., 2017], ssGSEA [Barbie et al., 2009] and scSE [Pont et al., 2019] within PLAID, using PLAID as ‘back-end’. The following functions have been developed within the PLAID framework: *replaid*.*sing, replaid*.*scse, replaid*.*ssgsea, replaid*.*gsva, replaid*.*ucell* and *replaid*.*aucell* (Supplemental data: “*Replicating other methods’ ESs*”). These functions are highly efficient in runtime and memory, generating ESs concordant with the original methods.

## 3 Results

Single-sample gene set and pathway scoring enable the prioritization of molecular signatures of individual samples, highlighting potential targets for personalized medicine. Several methods have been developed, each with distinct strengths and limitations. The most widely used include ssGSEA [Barbie et al., 2009], GSVA [Hänzelmann et al., 2013], AUCell [Aibar et al., 2017], singscore [Foroutan et al., 2018], scSE [Pont et al., 2019] and UCell [Andreatta et al., 2021]. These methods differ in preprocessing steps of ranking, centering, and normalization. Despite methodical variations, these methods collectively offer vast flexibility for data analysis [Fan et al., 2024]. Nevertheless, with the emergence of large-scale biological data (e.g., biobanks), most methods face substantial computational inefficiency. Except scSE, which is implemented in Go, other methods are in fact practically infeasible in runtime and memory requirements for large-scale data. This has limited the application of single-sample gene set scoring, hindering timely discovery.

We address this critical issue with PLAID (<u>P</u>athway <u>L</u>evel <u>A</u>verage Intensity <u>D</u>etection). Within each sample, PLAID identifies the genes mapped within a gene set and calculates the gene set score as the average log-intensity of genes in the gene set. PLAID does not zero-center or rank features, and leverages sparse matrices for efficient computations (Methods). We evaluated PLAID in single-cell transcriptomics (PBMC3K; Hao et al., 2023), bulk proteomics (Wolf et al., 2020), and microarray data (Lenz et al., 2008; https://www.cbioportal.org/datasets; Cerami et al., 2012]. These data span diverse scenarios in terms of resolution and distribution, thus allowing comprehensive benchmarks. We compared PLAID runtime, peak memory, and gene set scores to singscore [Foroutan et al., 2018], ssGSEA [Barbie et al., 2009], GSVA [Hänzelmann et al., 2013], scSE [Pont et al., 2019], UCell [Andreatta et al., 2021] and AUCell [Aibar et al., 2017]. Collectively these methods represent the most used single-sample gene set ES methods in biomedical research.

In scRNA-seq data, PLAID consistently outperformed other methods at different sample and gene sets sizes (Fig.1A; STable 1). PLAID completed scoring for 2864 gene sets in 1K cells in 0.17s, which is >100x faster compared to any other method. The second best method was scSE. Overall, the best memory-performing methods were PLAID and scSE. A similar trend was observed for 10K cells, where PLAID was >10x faster than other methods, while requiring similar RAM as other non-timed-out methods. When testing 61,459 gene sets in 1K cells, PLAID, singscore, UCell and AUCell successfully completed the gene set scoring. Other methods were timed out at 1h. PLAID completed the run in <8s, with <3GB peak RAM. Singscore, UCell and AUCell required >100x runtime compared to PLAID, with up to ∼1.5GB additional memory. GSVA, ssGSEA and scSE were timed out, and neither runtime nor peak RAM usage were accurately estimated (Fig.1A; STable 1). For 61,459 gene sets and 10K cells, PLAID achieved gene set scoring within 1h, requiring 110s and <20GB peak RAM (Fig.1A; STable 1). Other methods were timed out, and would have required at least 5x additional computing resources for a complete run.

While working with sparse matrices would be ideal, researchers often use dense matrices. For a comprehensive evaluation, we thus conducted runtime and memory profiling of PLAID on the TCGA-BRCA microarray dense data matrix [Cerami et al., 2012]. In line with previous observations, PLAID outperformed other methods in most scenarios. For instance, PLAID was >100x faster than AUCell and GSVA for 61459 gene sets and 1K cells, requiring 2.4GB. Altogether, these data demonstrate that PLAID is a highly efficient alternative to other single-sample gene set ES methods, but also that scSE remains a highly memory-optimized method capable of outperforming PLAID in some cases (SFig.1). We mapped runtime and peak RAM memory required by PLAID in detail at different combinations of gene sets and samples (SFig.2). Both runtime and peak memory usage increase approximately linearly with the number of cells or gene sets. For 1K gene sets and 1M cells, PLAID required about 200s and 28Gb RAM peak memory (SFig.2).

PLAID ESs are median normalized to aid cross-group comparisons (Methods). While conceptually based on a self-defined metric, we nevertheless compared PLAID with other methods. We found that in scRNA-seq, for the first available cell, PLAID ESs correlated well with singscore, UCell and AUCell ESs (Fig.1B). By enabling calculation of mean (Methods), scSE produces ESs highly concordant with PLAID (Fig.1B). Lower similarities emerged with GSVA and ssGSEA. Notably, GSVA and ssGSEA also exhibited low concordance with any other method (Fig.1B). Critically, within any single sample or cell, the raw gene set ESs are not well suited for direct comparisons between gene sets. Comparing gene sets would need statistical testing between samples, or centering gene sets across samples to obtain relative scores corresponding to gene set average centered log-expression. On the basis of this principle, we compared PLAID ESs vs other methods’ ESs after centering (Fig.1B). Interestingly, we observed a generalized improved concordance with all methods, including with GSVA and ssGSEA. This supports the suitability of PLAID ESs for differential testing between groups, and the possibility of cross-validate analyses from other methods. Testing in a bulk proteomics dataset (Wolf et al., 2020) confirmed high similarity between PLAID and scse.mean, and high concordance with singscore and ssGSEA (SFig.3). Low concordance emerged between PLAID and GSVA, in line with previous observations. Both in scRNA-seq and bulk proteomics, GSVA was lowly correlated with any other methods (SFig.3). Improved concordance was reached upon gene set centering -as per above-, supporting results in scRNA-seq data.

Running independent methods to validate enriched gene sets is a best practice. However, given the computational inefficiency of most current methods, this may be time-prohibitive in large datasets. We provide a solution to this problem by equipping PLAID with the most widely used single-sample gene set ES methods. We’ve implemented the following functions using PLAID as ‘back-end’: *replaid*.*sing, replaid*.*scse, replaid*.*ssgsea, replaid*.*gsva, replaid*.*ucell, replaid*.*aucell*. These functions provide efficient calculations of singscore, scSE, ssGSEA, GSVA, UCell and AUCell (Supplemental data, section “*Replicating other methods’ enrichment scores*”). To ensure accurate replication of the original methods, we conducted testing in scRNAseq, proteomics and microarray expression data (Fig.1C; SFig.4-9). Best concordances are seen for *replaid*.*sing, replaid*.*scse, replaid*.*ucell* and *replaid*.*aucell* vs. each respective original method, with further improvements made possible by method-specific parameters that reflect original implementations (Methods). A somewhat relatively lower concordance (especially for scRNA-seq data) emerges between *replaid*.*ssgsea* and *replaid*.*gsva* vs. original ssGSEA and GSVA, respectively. These methods were originally not intended for scRNA-seq. Lower concordance is also related to the ‘alpha’ and ‘tau’ parameters (Methods), due to approximations of rank weightings and ECDF in PLAID. Nevertheless, we achieve good concordance in nearly all cases, with the unmatched advantage of reaching up to ten-fold gain in computational efficiency. Altogether, these data demonstrate the multi-task power of PLAID in providing (i) its own single-sample gene set ES; (ii) unmatched computational efficiency; (iii) a framework for the most used single-sample ES methods, with much higher efficiency; (iv) evidence that if implemented expertly, the R language can be highly efficient.

Altogether, these analyses demonstrate that PLAID is a highly-performing and accurate method for single-sample gene set scoring. PLAID is ultrafast and memory efficient, generating gene set scores highly concordant with existing methods. PLAID can be up to 100x faster and requires significantly less memory-reaching up to 5-fold reduction in memory usage-when compared to any other method in any dataset.

## 4. Discussion

Scoring gene sets and pathways is vital for identifying coordinated molecular changes that may drive disease, especially when analyzed at the single-sample or single-cell level to enable personalized medicine. However, most current single-sample gene set scoring algorithms are not suitable for large datasets due to high memory requirements and slow runtimes. We introduce PLAID, an ultrafast, memory-efficient gene set scoring method developed in R using sparse matrices for optimized runtime and memory management. PLAID calculates gene set scores based on average intensity. PLAID demonstrates to be much faster than existing methods while maintaining high concordance in results. Uniquely, PLAID can also replicate scores from the most popular single-sample gene set scoring methods singscore, scSE, ssGSEA, GSVA, UCell, and AUCell, but with significantly improved efficiency. Thus, PLAID serves both as a standalone scoring tool and as a platform for running other gene set scoring methods on large datasets at substantially reduced computational demands.

## Supporting information

STable 1

## Supplementary data

Supplementary data are available online.

## Data availability

PLAID is implemented in R language for statistical computing. PLAID source code and package are freely available with no restriction on GitHub (https://github.com/bigomics/plaid). PLAID is also available on the Omics Playground platform (https://bigomics.ch/omics-playground).

