## Supplemental Data for "PLAID: ultrafast single-sample gene set enrichment scoring"

BigOmics Analytics, Via Serafino Balestra 12, Lugano 6900, Switzerland

#### Supplemental data

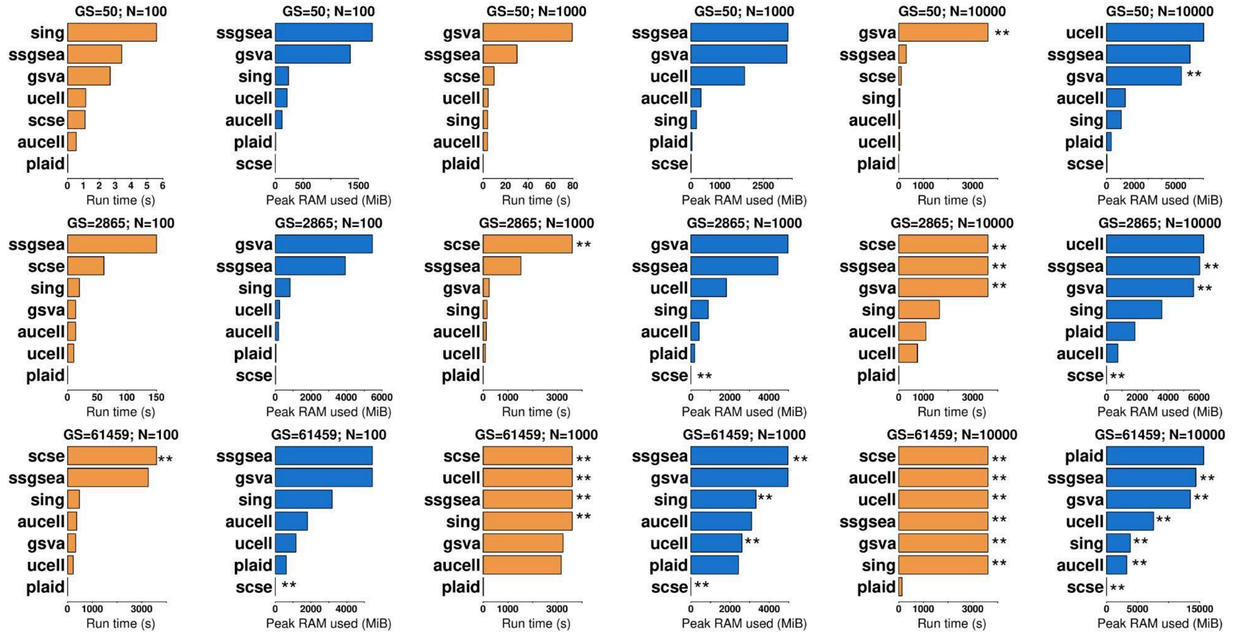


**Figure S1. Runtime and memory usage for all methods for dense dataset (TCGA-BRCA, RNA microarray; [Cerami et al., 2012]) with no sparsity**. Most methods are timed out after 3600 seconds (1 hour) for 61459 gene sets and 10K samples.

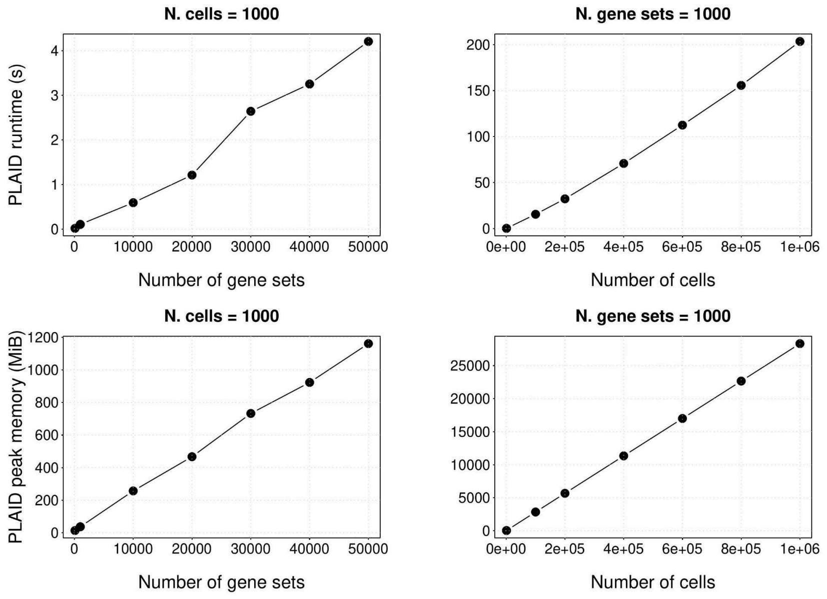


**Figure S2. PLAID runtime (s) and peak RAM memory (MiB).** Top and bottom left: representative testing of 1000 cells at an increasing number of gene sets (ranging from 10K to 50K). Top and bottom right: representative testing of 1000 gene sets at an increasing number of cells (ranging from 200K to 1M). Both runtime and peak memory usage increase approximately linearly with the number of cells or gene sets. For 1 million cells and 1000 gene sets, plaid took 203 seconds and required about 28Gb RAM of peak memory. All gene sets are part of the OmicsPlayground playbase gene set collection.

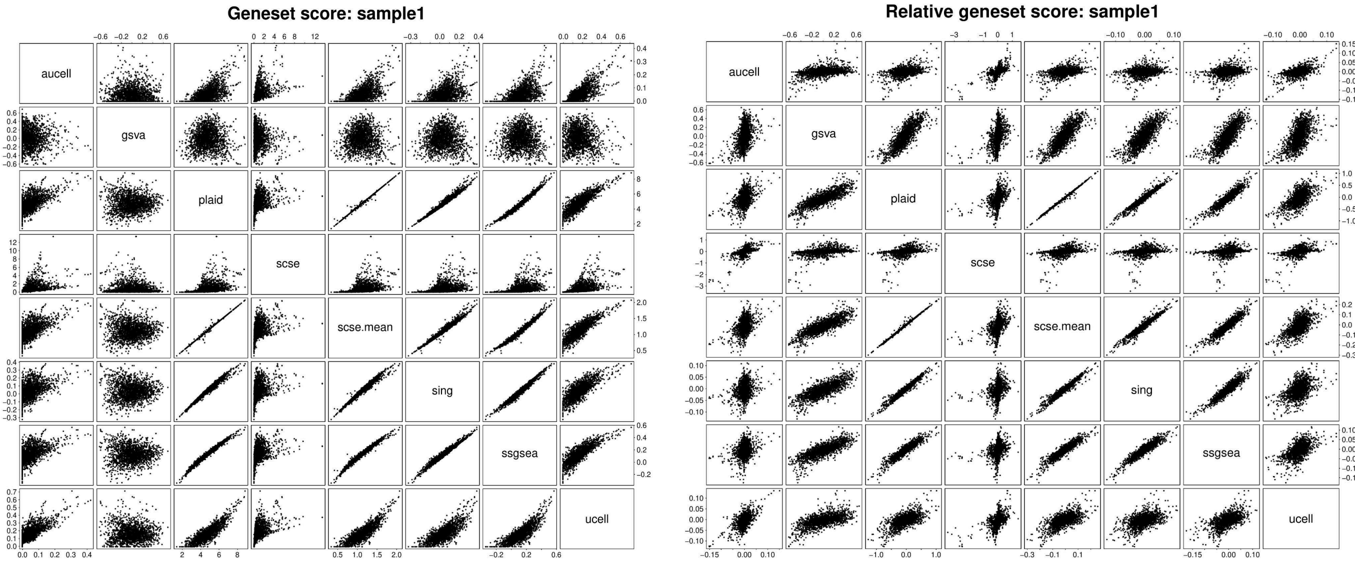


**Figure S3. Comparison between PLAID single-sample enrichment score and other methods' enrichment scores.** Pairwise scatter plots between PLAID's single-sample enrichment scores and single-sample enrichment scores generated by singscore, ssGSEA, GSVA, UCell, AUCell and scSE and scSE.mean (Methods). On the left, original enrichment scores. On the right, centered gene set enrichment scores. A bulk proteomics dataset (Wolf et al., 2020) was used as testing dataset. 2864 gene sets, corresponding to the OmicsPlayground collection of 'GO_BP' gene sets [Akhmedov et al., 2020], were used for analysis.

### Replicating other methods' enrichment scores using PLAID

#### Replication of singscore enrichment score with PLAID

Using the core PLAID function, we replicated the singscore enrichment score (Fouratan et al., 2018). The PLAID's singscore implementation is significantly faster. The following function *replaid.sing()*, which ranks and centers the input data matrix, is included in PLAID:

```
replaid.sing <- function(X, matG) {
  ## the ties.method=min is important for exact replication
  rX <- colranks(X, ties.method="min")
  rX <- rX / nrow(X) - 0.5
  gsetX <- plaid(rX, matG=matG, normalize=FALSE)
  return(gsetX)
}
```

Please refer to our GitHub repo at https://github.com/bigomics/plaid  to see the function in context.

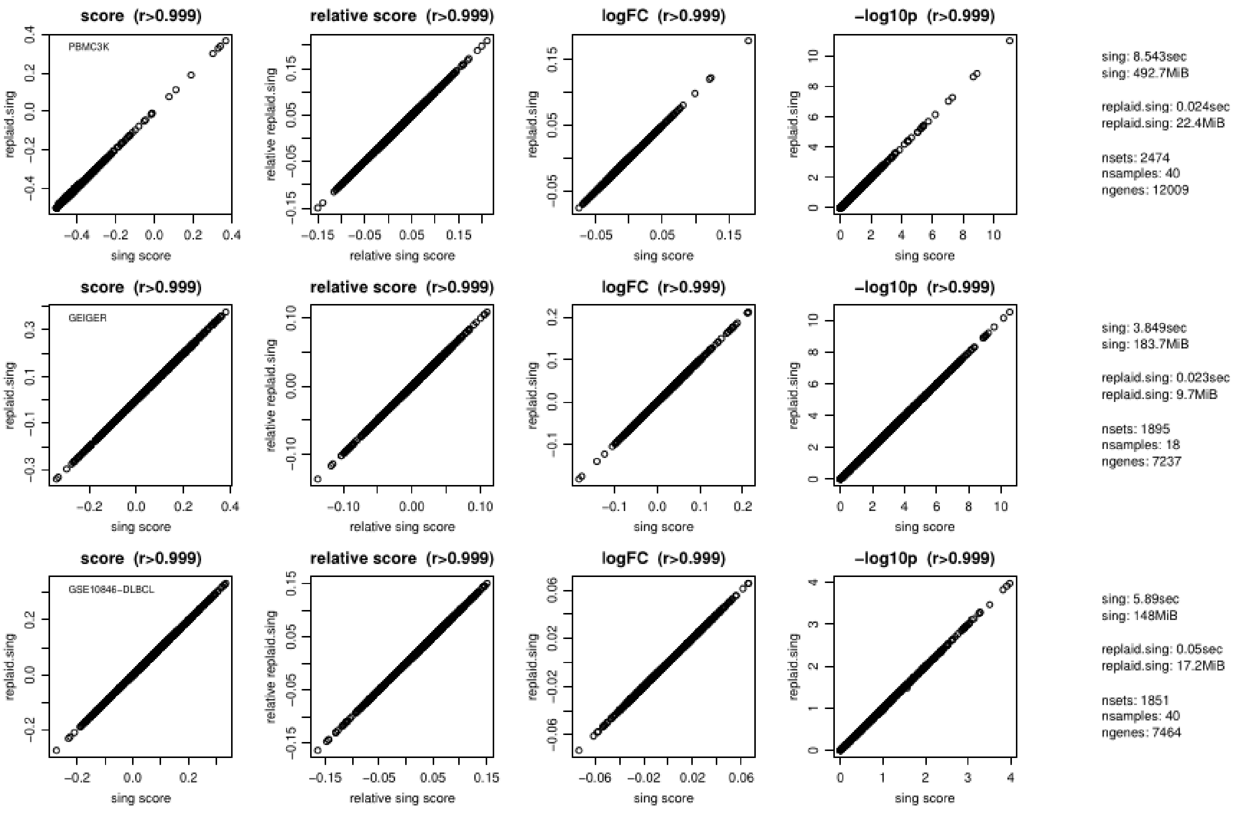


**Figure S4. Comparison between the original singscore algorithm (Fouratan et al., 2018) and the PLAID *replaid.sing()* R function.** Three testing datasets were used: (i) PBMC3K single-cell RNA-seq 10X Genomics dataset, including about 2700 single cells and freely available from the Seurat R package [Hao et al., 2023]; (ii) LC/MS proteomics dataset (Wolf et al., 2020); (iii) mRNA microarray (GSE10846; Lenz et al., 2008). Using a ranked expression matrix as input, the single-sample gene set ESs computed PLAID *replaid.sing(),* relative scores, the log2-fold-changes and differential gene expression p-values (on the -log10 scale) are perfectly correlated with singscore in all three tested datasets. Notably, the PLAID *replaid.sing()* is 10-50x faster than the original function and also requires less memory. We have confirmed similar results in other datasets (not shown).

#### Replication of scSE enrichment score with PLAID

Using the core PLAID function, we replicated the scSE enrichment score (Pont et al., 2019). The PLAID's scSE implementation is significantly faster. The following function *replaid.scse()*, which normalizes the output matrix with the total UMI intensity as is done by scSE, is included in PLAID:

```
replaid.scse <- function(X, matG, removeLog2=NULL, scoreMean=FALSE) {
  ...
  ## original scSE with Sum-statistics
  sX <- plaid(X, matG, stats="sum", normalize=FALSE)
  sumx <- Matrix::colSums(abs(X)) + 1e-8
  sX <- sX %*% Matrix::Diagonal(x = 1/sumx) * 100
  ...
}
```

Please refer to our GitHub repo at https://github.com/bigomics/plaid  to see the function in context.

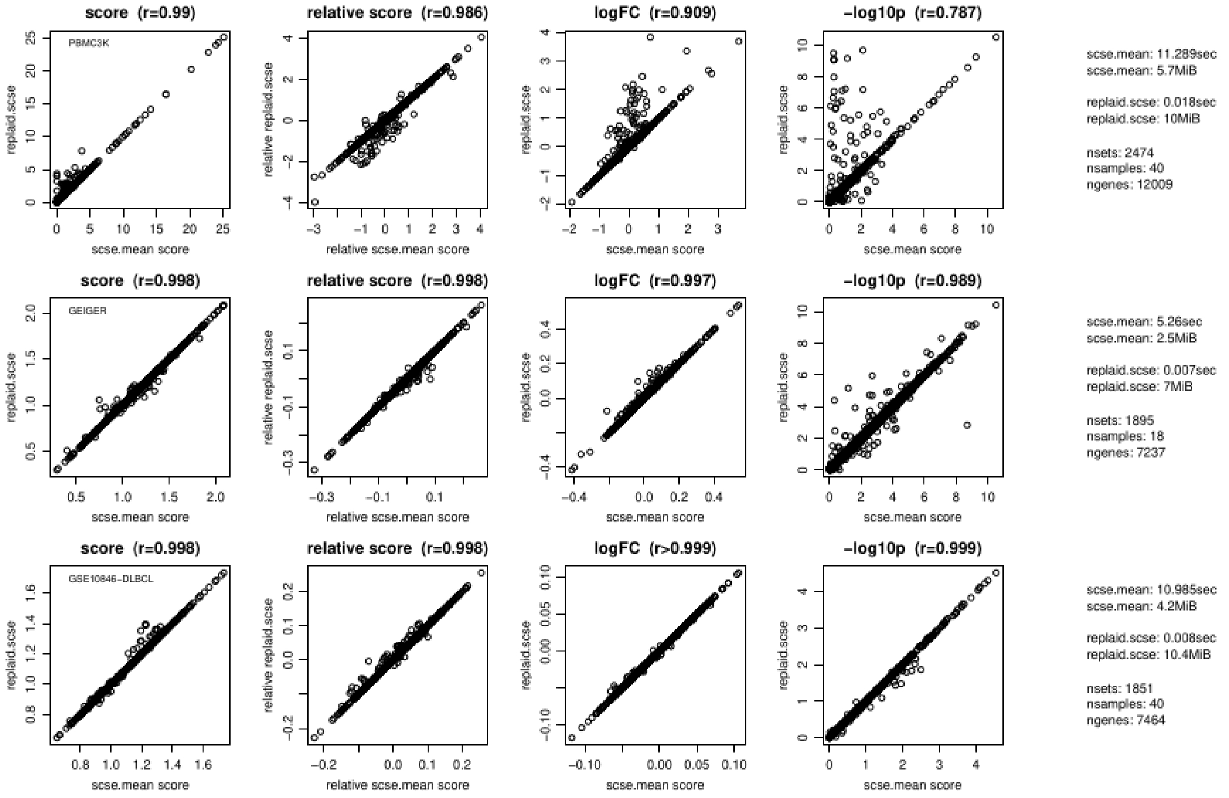


**Figure S5. Comparison between the original scSE algorithm (Pont et al., 2019) and the PLAID *replaid.scse()* R function**. Three testing datasets were used: (i) PBMC3K single-cell RNA-seq 10X Genomics dataset, including about 2700 single cells and freely available from the Seurat R package [Hao et al., 2023]; (ii) LC/MS proteomics dataset (Wolf et al., 2020); (iii) mRNA microarray (GSE10846; Lenz et al., 2008). The single-sample gene set enrichment scores computed PLAID *replaid.scse(),* relative scores, the log2-fold-changes and differential gene expression p-values (on the -log10 scale) are correlated with scSE in all three tested datasets. Correlation between the two methods can be enhanced by setting parameters '*removeLog2=FALSE*' and '*scoreMean=TRUE*' in *replaid.scse(). scoreMean*, when enabled, uses the mean of intensities rather than the sum of the intensities. This change in scSE, has also been discussed with the original author, Dr. F. Pont [Pont et al., 2019]. The PLAID *replaid.scse()* can be faster and require less memory than the original function. We have confirmed similar results in other datasets (not shown).

#### Replication of ssGSEA enrichment score with PLAID

Using the core PLAID function, we replicated the ssGSEA enrichment score (Barbie et al., 2009). The PLAID's ssGSEA implementation is significantly faster. The following function *replaid.ssgsea()*, which ranks and center the input data matrix, is included in PLAID:

```
replaid.ssgsea <- function(X, matG, alpha=0) {
  rX <- colranks(X, keep.zero=TRUE, ties.method="average")
  if(alpha != 0) rX <- rX^(1 + alpha)
  rX <- rX / max(rX) - 0.5
  dimnames(rX) <- dimnames(X)
  gsetX <- plaid(rX, matG, stats="mean", normalize=TRUE)
  return(gsetX)
}
```

For alpha>0 the values are less exact, nevertheless we tried to approximate the rank weighting in our code. Please refer to our GitHub repo at https://github.com/bigomics/plaid  to see the function in context.

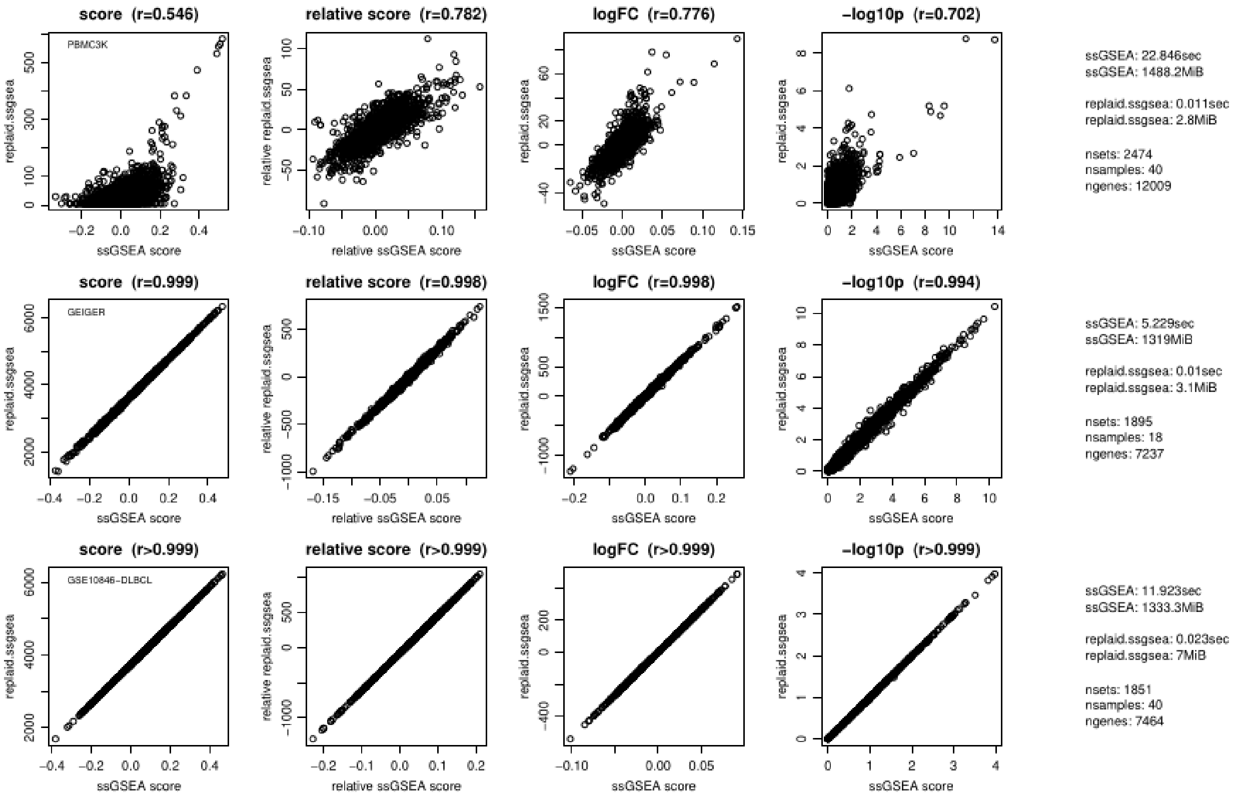


**Figure S6. Comparison between the original ssGSEA algorithm (Barbie et al., 2009) and the PLAID *replaid.ssgsea()* R function**. Three testing datasets were used: (i) PBMC3K single-cell RNA-seq 10X Genomics dataset, including about 2700 single cells and freely available from the Seurat R package [Hao et al., 2023]; (ii) LC/MS proteomics dataset (Wolf et al., 2020); (iii) mRNA microarray (GSE10846; Lenz et al., 2008). The single-sample gene set enrichment scores computed PLAID *replaid.ssgsea(),* relative scores, the log2-fold-changes and differential gene expression p-values (on the -log10 scale) are correlated with ssGSEA in all three tested datasets. Lower concordance is seen in the PBMC3K single-cell RNA-seq, presumably triggered by additional processing steps in ssGSEA not fully captured within PLAID and significantly affecting single-cell RNA-seq data. It is known that ssGSEA suffers from processing of single-cell RNA-seq data. Nevertheless, the concordance between PLAID *replaid.ssgsea()* results of plaid can be made highly similar to ssGSEA when using alpha=0 and unweighting the data. The PLAID's *replaid.ssgsea()* is much faster and requires less memory than the original function. We have confirmed similar results in other datasets (not shown).

#### Replication of GSVA enrichment score with PLAID

Using the core PLAID function, we replicated the GSVA enrichment score (Hänzelmann et al., 2013). The PLAID's GSVA implementation is significantly faster. The following function *replaid.gsva()*, which approximates the ECDF calculation, ranks and centers the input data matrix, is included in PLAID:

```
replaid.gsva <- function(X, matG, tau=0) {
  ## Faster approximation of relative activation
  zX <- (X - Matrix::rowMeans(X)) / (1e-8 + mat.rowsds(X))
  rX <- colranks(zX, signed=TRUE, ties.method="average")
  rX <- rX / max(abs(rX))
  if(tau > 0) rX <- sign(rX) * abs(rX)^(1 + tau)
  dimnames(rX) <- dimnames(X)
  ES <- plaid(rX, matG)
  return(ES)
}
```

For tau>0 the values are less exact even though we tried to approximate the rank weighting in our code. In the original formulation, GSVA uses an empirical CDF to transform expression of each feature to a [0;1] relative expression value. For efficiency reasons, this is approximated by a z-transform (center+scale) of each row, which is similar to the original implementation. Please refer to our GitHub repo at https://github.com/bigomics/plaid to see the function in context.

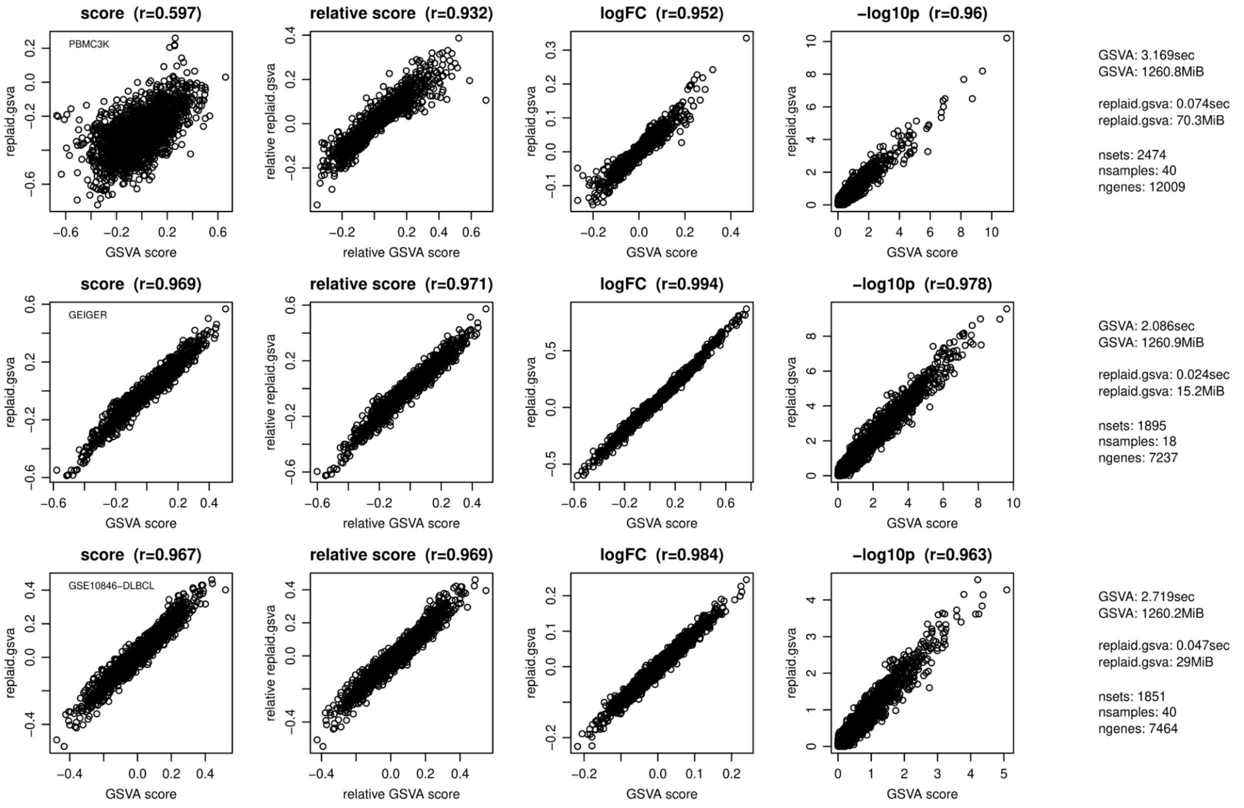


**Figure S7. Comparison between the original GSVA algorithm (Hänzelmann et al., 2013) and the PLAID *replaid.gsva()* R function**. Three testing datasets were used: (i) PBMC3K single-cell RNA-seq 10X Genomics dataset, including about 2700 single cells and freely available from the Seurat R package [Hao et al., 2023]; (ii) LC/MS proteomics dataset (Wolf et al., 2020); (iii) mRNA microarray (GSE10846; Lenz et al., 2008). The single-sample gene set enrichment scores computed PLAID *replaid.gsva(),* relative scores, the log2-fold-changes and differential gene expression p-values (on the -log10 scale) are correlated with GSVA in all three tested datasets. Lower concordance is seen in the PBMC3K single-cell RNA-seq, presumably triggered by additional processing steps in GSVA not fully captured within PLAID and significantly affecting single-cell RNA-seq data. Similarly to ssGSEA, GSVA also greatly suffers from processing of single-cell RNA-seq data. Nevertheless, the concordance between PLAID *replaid.gsva()* and original GSVA improves when using the relative score. The PLAID's *replaid.gsva()* is much faster and requires less memory than the original function. We have confirmed similar results in other datasets (not shown).

#### Replication of AUCell enrichment score with PLAID

Using the core PLAID function, we replicated the AUCell enrichment score (Aibar et al., 2017). The PLAID's AUCell implementation is significantly faster. The following function *replaid.aucell()*, which ranks and truncates the maximum rank as done in the original implementation, is included in PLAID:

```
replaid.aucell <- function(X, matG, aucMaxRank=ceiling(0.05*nrow(X))) {
  rX <- colranks(X, ties.method="average")
  ww <- 1.08*pmax((rX - (max(rX) - aucMaxRank)) / aucMaxRank, 0)
  gsetX <- plaid(ww, matG, stats="mean")
  return(gsetX)
 }
```

Please refer to our GitHub repo at https://github.com/bigomics/plaid  to see the function in context.

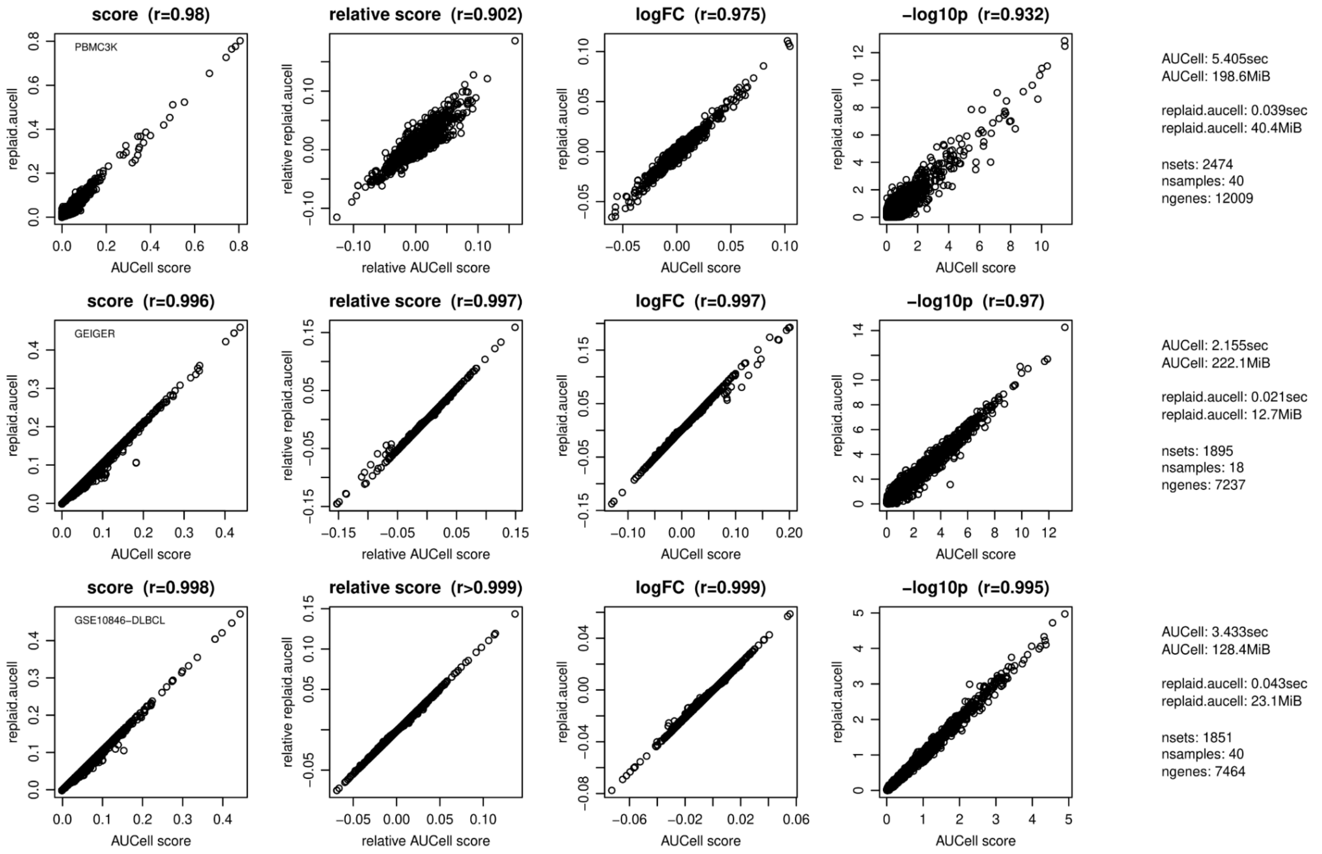


**Figure S8. Comparison between the original AUCell algorithm (Aibar et al., 2017) and the PLAID *replaid.aucell()* R function**. Three testing datasets were used: (i) PBMC3K single-cell RNA-seq 10X Genomics dataset, including about 2700 single cells and freely available from the Seurat R package [Hao et al., 2023]; (ii) LC/MS proteomics dataset (Wolf et al., 2020); (iii) mRNA microarray (GSE10846; Lenz et al., 2008). The single-sample gene set enrichment scores computed PLAID *replaid.aucell(),* relative scores, the log2-fold-changes and differential gene expression p-values (on the -log10 scale) are correlated with AUCell in all three tested datasets. The PLAID's *replaid.aucell()* is much faster and requires less memory than the original function. We have confirmed similar results in other datasets (not shown).

#### Replication of UCell enrichment score with PLAID

Using the core PLAID function, we replicated the UCell enrichment score (Andreatta et al., 2021). The PLAID's UCell implementation is significantly faster. The following function *replaid.ucell()*, which ranks and truncates the maximum rank as done in the original implementation, is included in PLAID:

```
replaid.ucell <- function(X, matG, rmax=1500) {
  rX <- colranks(X, ties.method="average")
  rX <- pmin(max(rX) - rX, rmax+1)
  S <- plaid(rX, matG)
  gsetX <- 1 - S / rmax + (Matrix::colSums(matG!=0)+1)/(2*rmax)
  return(gsetX)
}
```

Please refer to our GitHub repo at https://github.com/bigomics/plaid to see the function in context.

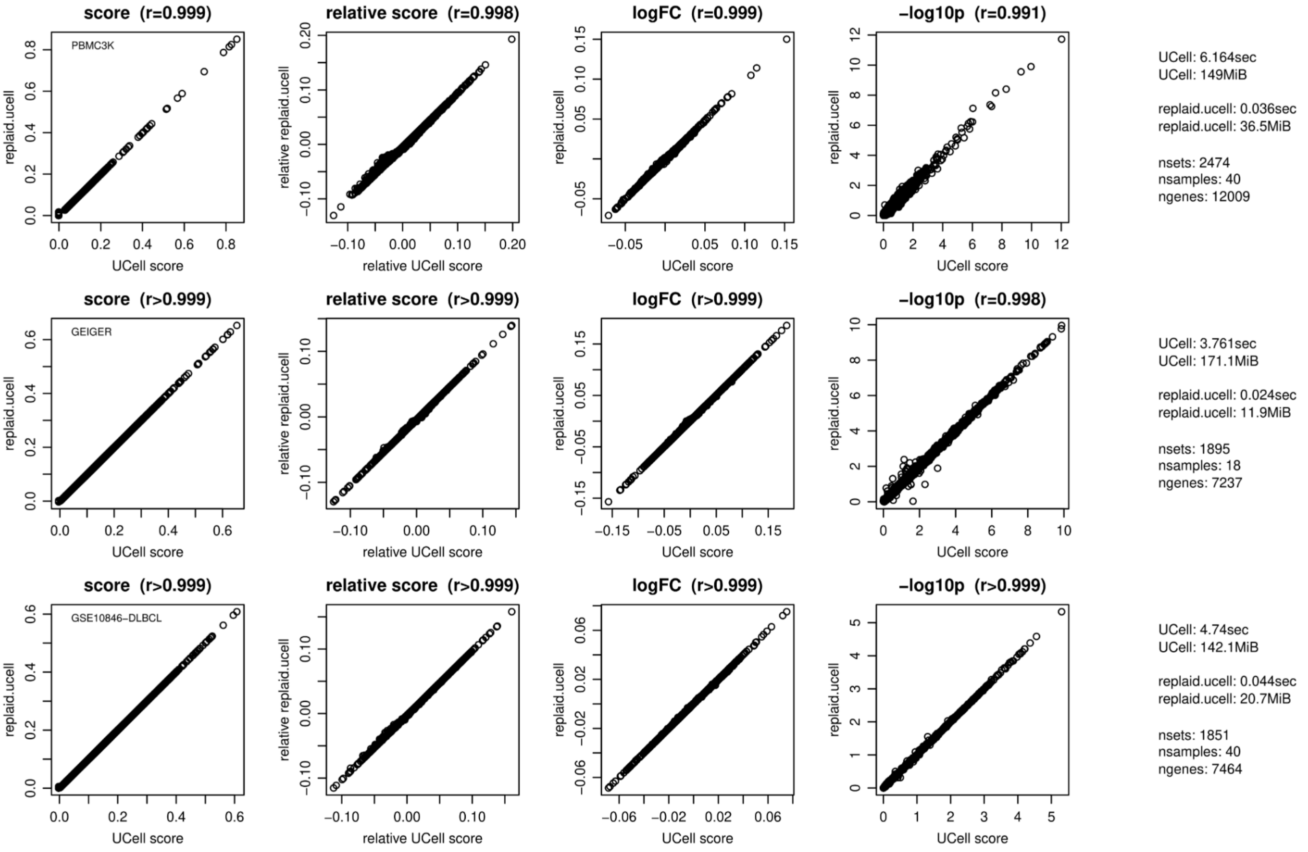


**Figure S9**. **Comparison between the original UCell algorithm (Andreatta et al., 2021) and the PLAID *replaid.ucell()* R function**. Three testing datasets were used: (i) PBMC3K single-cell RNA-seq 10X Genomics dataset, including about 2700 single cells and freely available from the Seurat R package [Hao et al., 2023]; (ii) LC/MS proteomics dataset (Wolf et al., 2020); (iii) mRNA microarray (GSE10846; Lenz et al., 2008). The single-sample gene set enrichment scores computed PLAID *replaid.ucell(),* relative scores, the log2-fold-changes and differential gene expression p-values (on the -log10 scale) are correlated with UCell in all three tested datasets. The PLAID's *replaid.ucell()* is much faster and requires less memory than the original function. We have confirmed similar results in other datasets (not shown).
